# Human Puf-A, a Novel Component of 90S Pre-ribosome, Links Ribosome Biogenesis to Cancer Progression

**DOI:** 10.1101/281857

**Authors:** Huan-Chieh Cho, Yenlin Huang, Jung-Tung Hung, Li-Chun Lai, Sheng-Hung Wang, Yun-Hen Liu, I-Ming Cho, Ming-Wei Kuo, Pei-Yi Cheng, Ming-Yi Ho, Rey-Jin Lin, Alice L. Yu, John Yu

**Author notes:** Address correspondence to John Yu, MD, PhD, No.5, Fuxing St., Guishan Dist., Taoyuan City 333, Taiwan, #5218; #5214.

## Abstract

We describe a novel biogenesis factor of the 90S pre-ribosome, Puf-A, which is a negative transcriptional target of p53. The expression of Puf-A is not only upregulated in advanced human lung cancer and tumors of patients especially with *TP53* mutation, but also is highly prognostic for stage I lung cancer. Loss of Puf-A expression prevents Kras^G12D^/p53^-/-^–induced tumor progression in the lungs and induces apoptosis in *TP53*– mutated cancers and c-Myc/p53^-/-^–transformed cells as well. Overexpression of Puf-A enhances proliferation of normal cells after c-Myc induction and overcomes the cell-cycle checkpoints incurred by p53 expression. Mechanistically, Puf-A interacts with double-stranded structures of the 5.8S sequence within pre-rRNA and maintains the integrity of 90S pre-ribosomes, thereby impacting early ribosome assembly and export of ribosomes from nuclei. Silencing of Puf-A disrupts the assembly of 90S pre-ribosomes and induces the translocation of its associated nucleophosmin (NPM1) from nucleoli to the nucleoplasm, resulting in impairment of ribosome synthesis. Thus, Puf-A is crucial for over-activation of ribosome biogenesis and contributes to tumor progression and cancer growth.

## Introduction

Oncogenic Kras activates both the Raf/MEK/ERK and PI3K/Akt/mTOR signaling pathways and subsequently enhances c-Myc expression, which is involved in cancer initiation and/or maintenance (McCubrey, Steelman et al., 2012). Mice with *KRAS* and *TP53* dual mutations produce phenotypically more advanced and faster growing lung tumors that are more resistant to chemotherapy compared to mice with *KRAS* mutation alone (Chen, Cheng et al., 2012, Oliver, Mercer et al., 2010). Clinically, both *KRAS* and *TP53* mutations in patients with lung cancer are important prognostic markers for shortened survival and have been associated with significantly worse outcomes for cytotoxic chemotherapy than observed with wild-type genotypes (Graziano, Gu et al., 2010, Tsao, Aviel-Ronen et al., 2007).

Enhancement of ribosome biogenesis activity is crucial for cell proliferation, transformation and consequent tumorigenesis through activation of *RAS* and *MYC* or inactivation of *TP53* (Truitt & Ruggero, 2016). During early ribosome biogenesis in the nucleolus, the long pre-rRNA is assembled with about 70 ribosome biogenesis factors and several small nucleolar RNAs (snoRNAs) to form a large 90S pre-ribosome, which is a critical precursor of 40S, 60S and 80S ribosomes (Oeffinger, 2016). Recently, the architecture of 90S pre-ribosome in yeast was unveiled and further revealed how the pre-rRNA folding, processing, and assembly at the first stage of ribosome biogenesis (Chaker-Margot, Barandun et al., 2017, Kornprobst, Turk et al., 2016, Sun, Zhu et al., 2017). Although many studies have shown various factors to be crucial for 90S assembly in yeast (Kressler, Hurt et al., 2017), it remains unclear which are the key factors involved in maintaining the integrity of 90S in mammalian cells. Therefore, unraveling the detailed machinery of 90S ribosome assembly could steer new therapeutic directions for *KRAS-* and/or *TP53*-mutated cancers.

Through comparative evolutionary genomic analysis, we first identified a novel protein, Puf-A (also known as KIAA0020 or PUM3), which belongs to a Puf-family of RNA-binding proteins highly expressed in primordial germ cells (Kuo, Wang et al., 2009). Puf-A plays an important role in the formation and specification of germ cell lineage, as knockdown of Puf-A specifically resulted in abnormal migration of primordial germ cells (Kuo et al., 2009). However, the pathophysiologic roles of Puf-A in cancer cells are largely unknown. The present study was undertaken to further identify the molecular machinery for regulation of Puf-A expression and determine its critical roles in ribosome biogenesis and cancer progression in order to provide both a diagnostic marker and a therapeutic target for combating *TP53*–mutated cancers.

## Results

### 1. Puf-A is upregulated in advanced human lung cancer and tumors with *TP53* mutation

The expression of Puf-A protein and RNA in human lung adenocarcinomas (ADCs) and squamous cell carcinomas (SCCs) was first examined and found to be elevated compared to non-neoplastic lung samples (Fig. 1A and Fig. S1a, left panel). According to a histological scoring system, H-score (Table S1), the expression of Puf-A was increased significantly in high grade tumors compared to low grade (Fig. 1A). Furthermore, grade 3 tumor cells exhibited not only greater Puf-A expression in nucleoli but also in the nucleoplasm (Fig. 1A) compared to grade 1 or 2 tumor cells. Also, *PUF-A* RNA was significantly increased in higher grade and advanced stage tumors (Fig. S1a, middle and right panels). The Kaplan-Meier analysis of public datasets revealed that increased *PUF-A* RNA expression correlated with shortened overall survival as well as progression-free survival of patients with lung cancer (Fig. 1B, left panel and Fig. S1b). Notably, the expression of Puf-A is highly prognostic for stage I lung cancer in our study. While the 5-years overall survival of stage I patients is 67.8%, consistent with the reported 50-70%(Der, Zhu et al., 2015), those with high and low Puf-A expression was 57.0% and 85.7%, (*p* = 0.038) respectively (Fig. 1B, right panel and Table S2). If confirmed in a larger study, Puf-A may be an important novel prognostic marker for stage I lung cancer.

**Figure 1.**
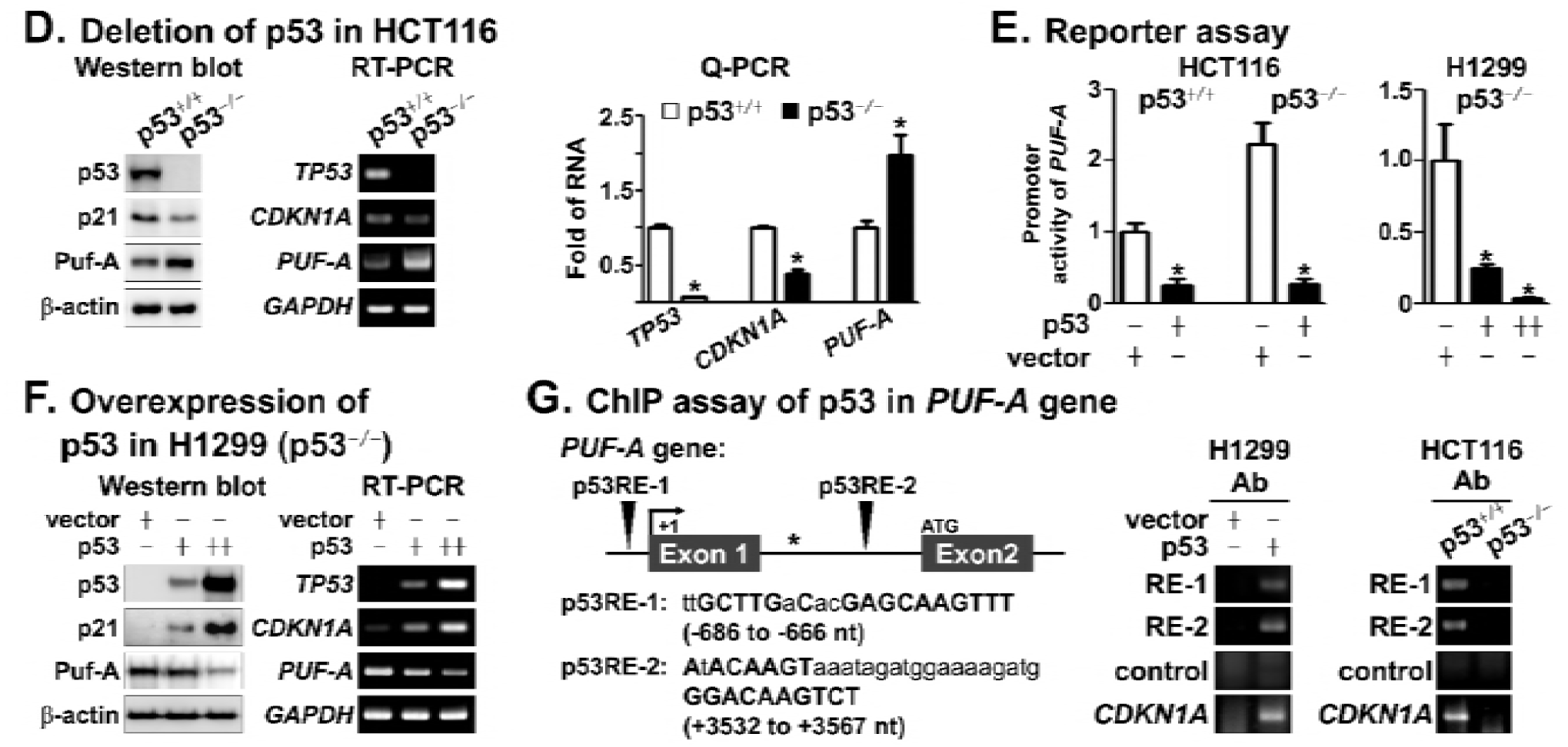
Expression of Puf-A in human lung cancer A. Immuno-histochemical analysis of tissue sections with different grades of lung cancer stained with antibody directed against Puf-A. The enlarged images of Puf-A staining in tissue sections of lung cancer are shown in the lower panels. Evaluation of Puf-A expression in various human lung cancers by a histological scoring system, H-score (Ishibashi, Suzuki et al., 2003), is shown in the right panel; Mean ± SEM values are indicated with red lines. ^**^ and ^***^ refer, respectively, to *p* < 0.01 and *p* < 0.001 (one-way ANOVA). **B**. Ten independently published RNA microarray datasets were analyzed using Kaplan Meier-plotter software (Gyorffy, Surowiak et al., 2013) as described in “Experimental Procedures”. Kaplan-Meier plots for first progression survival and overall survival for patients with Puf-A expression are shown (left panel). The Kaplan-Meier analysis of overall survival of patients with stage I lung cancer by evaluation of Puf-A expression. The high and low expression of Puf-A in stage I lung cancer was analyzed using “survival” and “survminer” packages in R version 3.4.1 (right panel). **C**. Immuno-histochemical analysis of p53 expression in tissue sections of lung cancer. The intensity of p53 staining was divided into two groups: “p53 mut” refers to tumor cells with strong nuclear p53 staining and “p53 WT” refers to those with weak or no staining. The percentages of p53 WT and mut tumor cells from patients with different grades of lung cancer is shown in the middle panels. Evaluation of Puf-A expression in tumor cells with p53 WT and mut by a histological scoring system, H-score, is shown in the right panel; Mean ± SEM values are indicated with red lines. ^**^ refers to *p* < 0.01 (two-tailed *t*-test). **D**. Western blot, RT-PCR and Q-PCR analyses of Puf-A (*PUF-A*), p53 (*TP53*) and p21 (*CDKN1A*) expression in p53^+/+^- and p53^-/-^-HCT116 cells were performed. β-actin and GAPDH were used, separately, as internal controls. In Q-PCR, Mean ± SD values (*n* = 3) are shown. ^*^ refers to *p* < 0.05 (one-way ANOVA). **E**. The promoter activity of *PUF-A* in p53^+/+^- and p53^-/-^-HCT116 cells after transfection with p53 was determined. Similarly, the promoter activity of *PUF-A* in p53^-/-^-H1299 cells after transfection with low (+) and high (++) amounts of p53 was also determined. Mean ± SD values (n = 3) are shown. ^*^ refers to *p* < 0.05 (one-way ANOVA). **F**. Western blot and RT-PCR analyses for Puf-A (*PUF-A*), p53 (*TP53*), and p21 (*CDKN1A*) expression in p53^-/-^-H1299 cells after transduction with control vector, and low and high titers of p53 virus were performed (*n* = 3). **G**. ChIP assay indicated the presence of two p53 response elements (p53RE-1 and p53RE-2, with the specific sequences shown) in the *PUF-A* gene (left panel). Antibody against p53 was used to immunoprecipitate the fragmented chromatin from H1299 cells after transduction with p53. Similarly, a ChIP assay of p53 protein that binds to regions of the *PUF-A* gene in p53^+/+^- and p53^-/-^-HCT116 cells was also performed (right panel). An equivalent amount of rabbit IgG was added as the negative control. p53RE-1, p53RE-2 and control DNA (denoted with ^*^) sequences of *PUF-A* regions or p53RE of *CDKN1A* regions were analyzed by semi-quantitative PCR.

It should be noted that *PUF-A* was significantly increased in tumor cells of patients with mutated *TP53* mutation (Fig. S1c) compared to tumors with wild-type genotypes. To assess whether the status of *TP53* correlates with Puf-A expression, tissue sections from patients with lung cancer were further evaluated for the expression of p53 by immuno-histochemical analysis (Fig. 1C and Table S1). Strong p53 nuclear staining was detected in 31 (37.8%) tumor samples from patients with lung cancer (Fig. 1C, left panel; p53 mut), suggesting that these tumors frequently harbored mutated *TP53* (Xie, Lan et al., 2014). Other samples with weak or no staining of p53 conceivably had wild-type p53 which has a short half-life (Fig. 1C, left panel; p53 WT). Interestingly, the percentage of samples with high p53 expression in these patients was correlated with tumor grade (Fig. 1C, middle panel). Moreover, H-score analysis showed that Puf-A expression was significantly increased in tumors with high p53 expression, compared to those with low p53 (Fig. 1C, right panel), suggesting that p53 could regulate Puf-A expression.

In p53^+/+^-H460 (lung cancer) and -HCT116 (colon cancer) cells after silencing of p53 expression, the expression of Puf-A protein and RNA was significantly increased (Fig. S1d), consistent with findings of higher Puf-A expression in p53^-/-^- as compared to p53^+/+^-HCT116 cells (Fig. 1D). In contrast, exogenous p53 significantly suppressed the promoter activity of *PUF-A* in both p53^+/+^- and p53^-/-^-HCT116 cells (Fig. 1E, left panel). Similarly, in p53^-/-^-H1299 (lung cancer) cells, exogenous p53 not only reduced the promoter activity of *PUF-A* (Fig. 1E, right panel) but also Puf-A protein and RNA expression in a dose-dependent manner compared to control cells (Fig. 1F). Furthermore, the chromatin immunoprecipitation (ChIP) assay revealed that p53 binds directly to the p53RE-1 and -2 sites of *PUF-A* locus in H1299 cells after transduction with exogenous p53 or in p53^+/+^-HCT116 cells (Fig. 1G). These results indicate that p53 transcriptionally suppresses Puf-A expression.

### 2. Puf-A is transcriptionally regulated by oncogenic Kras^G12D^ and c-Myc

Oncogenic Kras signaling was reported to be associated with c-Myc expression in cancer (DeNicola, Karreth et al., 2011). The *Puf-A* expression at the RNA level was significantly increased in tumors of patients with mutated *KRAS* and high *MYC* expression compared to tumors with wild-type genotypes (Fig. S1e). In H1299 cells after transduction with Kras^G12D^ or c-Myc, the expression levels of Puf-A protein and RNA were increased, respectively, compared to control cells (Fig. 2A and 2B). Furthermore, both Kras^G12D^ and c-Myc increased *PUF-A* promoter activity by 2.2- and 3.2-fold (Fig. 2C). Moreover, the ChIP assay showed that the exogenous c-Myc protein bound the c-Myc response element (c-MycRE) in exon 1 of the *PUF-A* gene but not the control DNA sequence (Fig. 2D, left panel). Similarly, the endogenous c-Myc protein also bound the c-MycRE in exon 1 of the *PUF-A* gene in both p53^+/+^- and p53^-/-^-HCT116 cells (right panel). These results indicated that c-Myc transcriptionally activated the expression of *PUF-A*.

**Figure 2.**
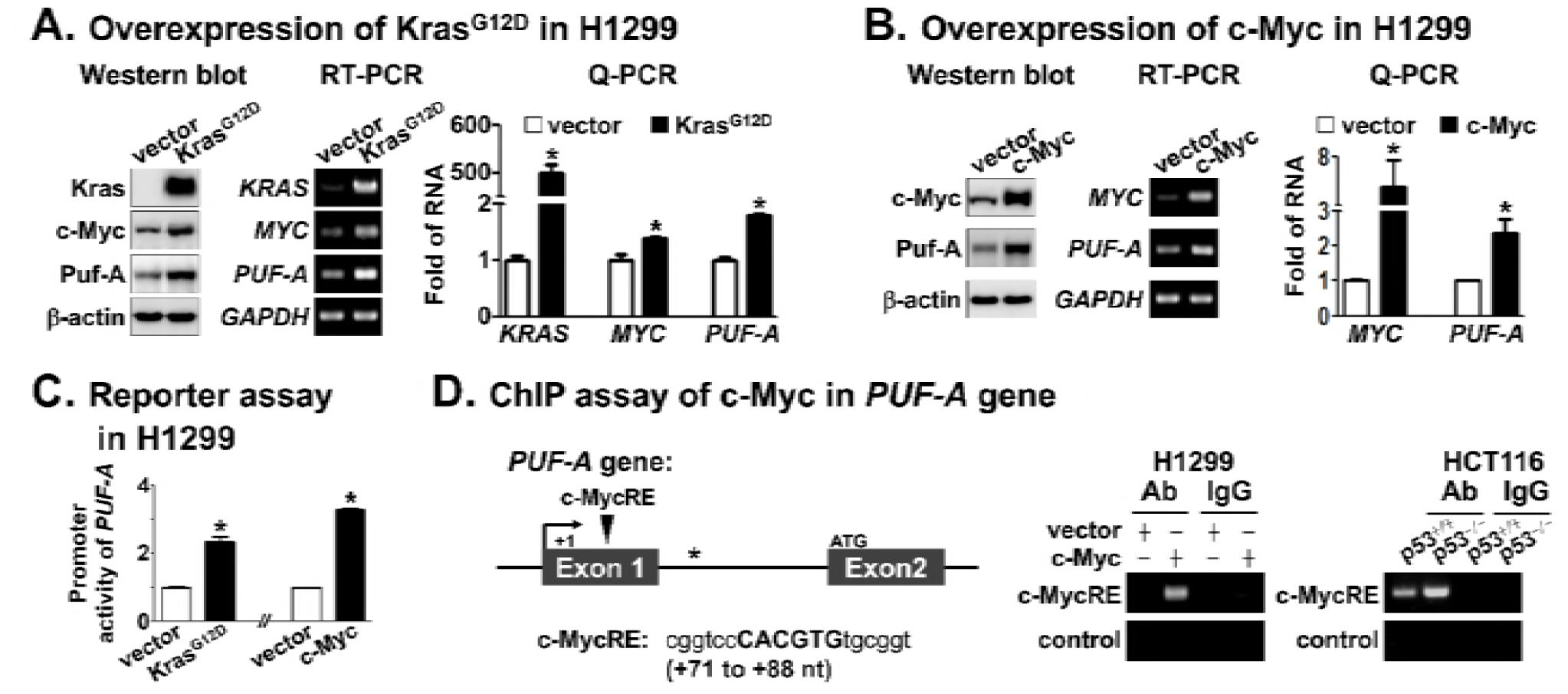
Transcriptional regulation of Puf-A expression and inhibition of tumor progression in the lungs through Puf-A silencing A. Western blot, RT-PCR and Q-PCR analyses of Kras (*KRAS*), c-Myc (*MYC*) and Puf-A (*Puf-A*) expression in Kras^+/+^-H1299 cells after transduction with either control vector or Kras^G12D^ were performed. β-actin and GAPDH were used, separately, as internal controls. In Q-PCR, Mean ± SD values (*n* = 3) are shown. ^*^ refers to *p* < 0.05 (one-way ANOVA). **B**. Western blot, RT-PCR and Q-PCR analyses of c-Myc (*MYC*) and Puf-A (*Puf-A*) expression in Kras^+/+^-H1299 cells after transduction with either control or c-Myc overexpression vectors were performed. β-actin and GAPDH were used, separately, as internal controls. In Q-PCR, Mean ± SD values (*n* = 3) are shown. ^*^ refers to *p* < 0.05 (one-way ANOVA). **C**. The promoter activity of *PUF-A* in H1299 cells after transfection with control and Kras^G12D^ or c-Myc vectors. Mean ± SD values (n = 3) are shown. ^*^ refers to *p* < 0.05 (one-way ANOVA). **D**. ChIP assay indicated the presence of c-Myc response element (c-MycRE, with the specific sequences shown) in the *PUF-A* gene (left panel). Antibody against c-Myc was used to immunoprecipitate the fragmented chromatin from H1299 cells after transduction with c-Myc. Similarly, a ChIP assay of c-Myc protein that binds to regions of the *PUF-A* gene in p53^+/+^- and p53^-/-^-HCT116 cells was also performed (right panel). An equivalent amount of rabbit IgG was added as the negative control. c-MycRE and control DNA (denoted with ^*^) sequences of *PUF-A* regions were analyzed by semi-quantitative PCR. **E**. Lung tissues with tumors from CCSP-rtTA/TetO-Cre/LSL-Kras^G12D^/p53^+/+^ mice at 6 and 12 weeks after administration of doxycycline for Kras^G12D^ activation were stained with antibodies directed against Puf-A. The bronchiolar and alveolar regions of lung lesions are shown; enlarged images of Puf-A staining in the enclosed areas are shown in the lower panels. **F**. Lung tissues from CCSP-rtTA/TetO-Cre/LSL-Kras^G12D^/p53^flox/flox^ mice at 2 weeks and 8 weeks after administration of doxycycline for Kras^G12D^ activation and p53 deletion were stained with antibodies directed against Puf-A. The bronchiolar and alveolar regions of the lung sections are shown; enlarged images of Puf-A staining in the enclosed areas are shown in the lower panels. **G**. The KP-1 and KP-2 mouse lung adenocarcinoma cell lines derived from CCSP-rtTA/TetO-Cre/LSL-Kras^G12D^/p53^-/-^ mice after administration of doxycycline for eight weeks were examined for the silencing efficiency of Puf-A expression. Western blot and Q-PCR analyses for Puf-A expression in KP-1 and KP-2 cells after transduction with control shLacZ, shPuf-A-1, and shPuf-A-2 are shown (shRNA sequences for silencing mouse Puf-A are listed in Table S2). **H**. The experimental design after *in vivo* intranasal administration of lentivirus to silence Puf-A expression in the lung tissues of CCSP-rtTA/TetO-Cre /LSL-Kras^G12D^/p53^flox/flox^ mice after simultaneous Kras^G12D^ activation and p53 deletion. **I**. The decrease in the number of tumor foci in mice after intranasal administration of, shPuf-A-1 counted in the tissue sections. Mean ± SEM values are shown for control shLacZ (*n* = 13), shPuf-A-1 (*n* = 11), and shPuf-A-2 (*n* = 6). ^*^ and ^***^ refer, respectively, to *p* < 0.05 and *p* < 0.001 (one-way ANOVA).

### 3. Silencing of Puf-A inhibits Kras^G12D^/p53^-/-^–induced tumor progression in lungs

We previously established an inducible lung ADC model in CCSP-rtTA/ TetO- Cre/LSL-Kras^G12D^/p53^+/+^ mice (Fig. 2E) that gave rise to adenoma and ADC in the lungs after Kras^G12D^ induction for 6 and 12 weeks, respectively (Cho, Lai et al., 2011). It was found that the Puf-A-positive cells were further increased in bronchiolar ADC of the lungs, while there were only a few Puf-A-positive cells scattered within the alveolar tumor region. Mutation of *KRAS* and *TP53* genes often occurs concurrently and with similar frequencies in human lung cancer (Yamaguchi, Kugawa et al., 2012). The progression from adenoma to ADC in mutant *KRAS*–driven mouse models can be accelerated through loss of the *TP53* tumor-suppressor gene (Jackson, Olive et al., 2005). In CCSP-rtTA/TetO-Cre/LSL-Kras^G12D^/p53^flox/flox^ mice, the first appearance of adenomas in the lung occurred at 2 weeks, and the progression from adenoma to ADC was accelerated at 8 weeks after Kras^G12D^ activation and p53 deletion (Fig. 2F). The intensity of Puf-A staining in ADC cells of bronchiolar and alveolar regions were greatly increased (Fig. 2F) compared to tumor cells in Kras^G12D^/p53^+/+^ mice after 12 weeks of activation (Fig. 2E), in agreement with the staining patterns of Puf-A in human lung cancer with *TP53* mutation (Fig. 1C).

Two ADC cell lines (KP-1 and KP-2) derived from Kras^G12D^/p53^-/-^–induced lung ADCs were used for selection of shRNAs for Puf-A silencing (Fig. 2G). After transduction with shPuf-A-1 and -2, the expressions of Puf-A decreased significantly in both lines of cells (Fig. 2G) compared with control cells. We further determined whether a reduction in Puf-A expression after the onset of adenomas might prevent progression to ADCs in the lungs. As shown in Figure 2D, the lentiviral vectors expressing shPuf-A-1, shPuf-A-2 or control shLacZ were delivered intranasally four times between weeks 2 and 4 to the lungs of CCSP-rtTA/TetO-Cre/ LSL-Kras^G12D^/p53^flox/flox^ mice. The tumor foci in the lungs were then examined after Kras^G12D^ activation and p53 deletion at week 8 (Fig. 2H). In the lungs treated with shPuf-A-1 or shPuf-A-2 lentivirus, adenoma formation was reduced by 1.9- and 1.5-fold, respectively, and adenocarcinoma formation was reduced by 4.0- and 2.6-fold, respectively, (Fig. 2I) compared to controls. These results support an essential role of Puf-A in the development and progression of ADC in the lungs.

### 4. Puf-A is essential for cancer growth and cell transformation

Various cancer cells, including p53-proficient (p53^+/+^-A549, p53^+/+^-H460 and p53^+/+^- HCT116 cells) and p53-deficient cells (p53^-/-^-HCT116, p53^-/-^-H1299, p53^R^248^W^-CL1-5 and p53^R^280^K^-MB231 cells), were used in studies on the silencing of Puf-A expression. After transduction with shPuf-A-1 and shPuf-A-2, the expression of Puf-A was markedly reduced compared to control cells (Fig. 3A). When Puf-A was silenced in p53-proficient cancers, the expression levels of p53 and p21 proteins increased greatly (Fig. S2a and S2b) compared with control cells. However, there was no apparent change in the expression level of cleaved caspase3 or cleaved poly ADP-ribose polymerase 1 (PARP1) in response to Puf-A silencing in p53-proficient cells (Fig. 3A, left panel). In contrast, after Puf-A silencing in the p53-deficient cells cleaved caspase3 and cleaved PARP1 were markedly increased (Fig. 3A, right panel).

**Figure 3.**
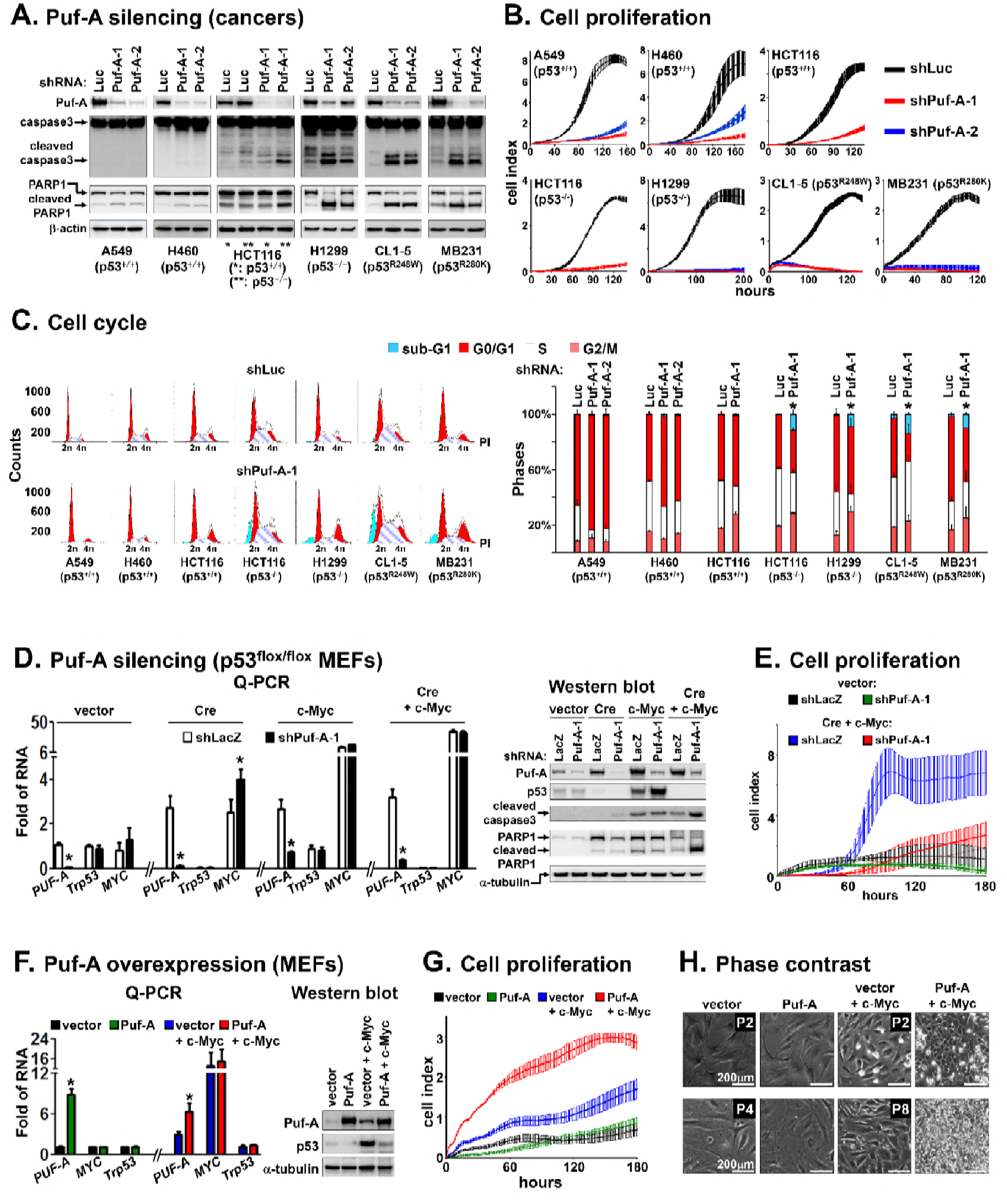
The effect of Puf-A on cell growth. A. Lentiviruses with two short hairpin RNAs (shPuf-A-1 and shPuf-A-2 sequences for silencing mouse Puf-A are listed in Table S2) were transduced into various cancer cell lines. The expression levels of Puf-A were greatly decreased in all cell lines after knockdown of Puf-A expression with shRNAs compared with cells (controls) transduced with shLuc. The cleaved forms of caspase3 and PARP1 increased significantly in p53^-/-^ cells after transduction with shPuf-A-1 and shPuf-A-2 lentivirus when compared to p53^+/+^ cells. In p53^+/+^ cells, silencing of Puf-A did not result in any apparent changes in the cleaved form of caspase3 and PARP1 when compared to control shLuc transduced cells. **B**. Proliferation of various cancer cell lines over time, including p53^+/+^-A549, -H460 and -HCT116 cells; p53^-/-^-HCT116 and -H1299 cells; p53^R^248^W^-CL1-5 cells and p53^R^280^K^- MB231 cells, after transduction with control shLuc, shPuf-A-1 and shPuf-A-2. **C**. Flow cytometric analysis of cell cycle distribution of Puf-A-silenced cells (same lines as in Fig. 3B). All cell lines were transduced with control Luc, shPuf-A-1, or shPuf-A-2 and then incubated with propidium iodide (PI) for assessment of cell cycle distribution based on DNA content. The sub-G1 cell population (blue) in p53^-/-^-H1299 and -HCT116 cells, p53^R^248^W^-CL1-5 cells and p53^R^280^K^-MB231 cells was significantly increased after transduction with shPuf-A-1 and shPuf-A-2 virus compared to shLuc transduced controls. Mean ± SD values (*n* = 3) are shown (one-way ANOVA). **D**. Q-PCR and western blot analyses of the expression of *PUF-A-, Trp53*- and *MYC*- RNA and Puf-A-, p53- and PARP1-protein in p53^flox/flox^ MEFs that either overexpressed Cre or c-Myc or co-expressed Cre with c-Myc. Transduction with vector control and shLacZ were used for comparison. Mean ± SD values (*n* = 3) are shown. ^*^ refers to *p* < 0.05 (one-way ANOVA). **E**. Proliferation of various cells over time, including MEFs transduced with control vector or Cre plus c-Myc, after transduction with control shLacZ and shPuf-A-1. **F**. Q-PCR and western blot analyses of the expression of *PUF-A-, MYC*- and *Trp53*- RNA, and Puf-A- and p53-protein in MEFs that either overexpressed Puf-A alone or co-expressed Puf-A and c-Myc. Transduction with vector control was used for comparison. β-actin and GAPDH were used, separately, as internal controls. Mean ± SD values (*n* = 3) are shown. ^*^ refers to *p* < 0.05 (one-way ANOVA). **G**. A proliferation assay was performed with MEFs that overexpressed c-Myc or co-expressed Puf-A with c-Myc. **H**. Phase contrast images of MEFs after transduction as described above in early- (upper panel, P2) and late-passages (lower panel, P4 and P8). P indicates the generation of passages after virus infection.

The proliferation rates of these cancer cells after silencing Puf-A was obviously reduced (Fig. 3B). The proportion of p53-proficient cells in the G0/G1 phase after Puf-A silencing increased greatly, whereas the proportion in the S phase was reduced (Fig. 3C, left panel), indicating that silencing Puf-A in p53-proficient cancers induces cell-cycle arrest. On the other hand, the proportion of p53-deficient cancer cells in the sub-G1 phase after silencing Puf-A significantly increased (Fig. 3C, right panel, blue color).

Furthermore, when Puf-A was silenced in p53^flox/flox^ mouse embryo fibroblasts (MEFs) co-expressing c-Myc and Cre (a recombinase for p53 deletion), the expression of cleaved caspase3 and cleaved PARP increased markedly (Fig. 3D). Also, Puf-A silencing obviously decreased the rate of proliferation of these cells (Fig. 3E) compared to cells transduced with control vector. These results indicate that silencing of Puf-A in various *TP53*–mutated cancers and c-Myc/p53^-/-^–transformed cells leads to apoptosis.

In early-passage (P2) MEFs with p53^+/+^, the expression of p53 was obviously suppressed after co-transduction with Puf-A and c-Myc (Fig. 3F and Fig. S2c, left panel) when compared to those transduced with vector and c-Myc. Furthermore, such co-transduction also increased the proliferation rate by 2.0-fold and greatly increased the percentage of Ki-67^+^ (a mitotic marker) cells (Fig. 3G and Fig. S2c, right panel). In late-passage (P8) MEFs with Puf-A and c-Myc co-transduction, cells grew to a much higher density and exhibited obvious phenotypic conversions (e.g., shrinkage of cell size, large nuclear-cytoplasmic ratio) than MEFs with c-Myc transduction alone (Fig. 3H). These results indicate that Puf-A enhances the ability of c-Myc to stimulate cell proliferation and enables cells to evade p53-mediated cell cycle arrest induced by c-Myc activation (Nilsson & Cleveland, 2003).

### 5. Puf-A is required for ribosome biogenesis

Increased rRNA transcription in nucleolus is a common feature of cancer cells, which require elevated ribosome biosynthesis to support growth (Baker, Cooke et al., 2017). We next examined whether Puf-A is involved in rRNA transcription, thereby affecting rRNA expression patterns. The steady state of pre-45S (a component of 90S pre-ribosome), 18S (a component of 40S ribosome), 5.8S and 28S (the components of 60S ribosome) rRNA expression in MEFs, p53^+/+^-HCT116, p53^-/-^-HCT116 and p53^-/-^- H1299 cells after various transductions to increase or silence Puf-A expression was analyzed. As shown in Fig. 4A, the results showed that there was not much difference in the expression level of these rRNAs, except for slight changes of pre-45S, in response to Puf-A overexpression or silencing.

Polysome profiling assays showed that the amount of 80S ribosome in MEFs after co-transduction with Puf-A and c-Myc increased remarkably when compared to cells transduced with c-Myc alone (Fig. 4B, left panel). In contrast, the amount of 80S ribosome in cancer cells (p53^+/+^-HCT116, p53^-/-^-HCT116 and p53^-/-^-H1299 cells), especially those with defective p53, decreased markedly after transduction with shPuf-A- 1 compared to control shLuc cells (Fig. 4B, right panels).

Furthermore, in ribosomal fractions of polysome profiling using MEFs and cancer cells, the protein distribution of S6 (RPS6, a marker for the 40S ribosome) and L5 (RPL5, a marker for the 60S ribosome) is shown in Figure 4C. If MEFs were co-transduced with Puf-A and c-Myc, the expression of S6 protein in 40S and 80S fractions and L5 protein in 60S and 80S fractions increased greatly (Fig. 4C II) compared to MEFs transduced with c-Myc alone (Fig. 4C I). However, in cancer cells silenced with shPuf-A-1, the expression of S6 in the 40S and 80S fractions and L5 in the 60S and 80S fractions decreased significantly (Fig. 4C IV, VI and VIII) compared to control shLuc cells (Fig. 4C III, V and VII). These results implied that Puf-A is involved in ribosome assembly.

### 6. Puf-A interacts with the 5.8S sequence within pre-rRNA of 90S pre-ribosomes

Western blotting of gradient fractions in polysome profiling assays for nuclear and cytosolic preparations indicated that the 40S, 60S, and 90S pre-ribosomes were found in the nuclear fraction (red curve), while the mature 40S, 60S, and 80S ribosomes and polysomes were in the cytosolic fraction (black curve) in p53^+/+^- and p53^-/-^-HCT116 cells (Fig. 5A, upper panel). Notably, protein distributions further showed that Puf-A was found primarily with nuclear 90S pre-ribosomes but not with cytosolic 80S ribosomes; in contrast, S6 and L5 were expressed in both the nucleus and cytoplasm (Fig. 5A, lower panel).

The sub-cellular localization of Puf-A in cells was further examined with specific antibodies in immunostaining. The result showed that HA-tagged Puf-A (red) was co-localized with pre-rRNA (stained with green 5.8S rRNA sequence antibody) primarily in the nucleus, particularly in the nucleolus, but not in the cytoplasm of H1299 cells (Fig. 5B), consistent with the staining profiles of Puf-A in lung tumor specimens (Fig. 1A; Fig. 2E and 2F). These analyses suggested that Puf-A specifically associates with 90S pre-ribosomes in the nucleolus.

Puf-A belongs to the PUF-family that has RNA-binding activity in variety of spices (Quenault, Lithgow et al., 2011). To examine whether Puf-A has the capacity to bind to particular pre-rRNA sequences of 90S pre-ribosomes in the nucleolus, purified recombinant human Puf-A protein (rPuf-A) was incubated with total RNA extracts from cells for a pull-down assay. There was a strong RNA band corresponding to a 100˜200 nt fragment in the agarose gel analysis (Fig. S3a, left panel). With the use of 5.8S sequence-specific primers, it was confirmed that this RNA fragment contained 5.8S sequence (Fig. S3a, right panel). Moreover, RNA-EMSA analyses showed that rPuf-A interacted with the nt 1-156, 37-156 and 37-103 sequences of 5.8S rRNA (Fig. 5C), but not the nt 1-40 and 103-156 sequences. In addition, rPuf-A was shown not to interact with 5S rRNA (Fig. S3b).

The interactions between Puf-A and the 5.8S sequences in the pre-rRNA were further examined by RNA-protein docking as described in Supplementary Experimental Procedures using NPDock server(Tuszynska, Magnus et al., 2015). Through examination of 100 potential models of Puf-A protein in complex with 5.8S rRNA sequences by NPDock, the model which possessed the lowest energy scores was considered as the predicted binding mode and shown in Fig. 5D and S3c. The 5.8S rRNA sequence in the model has 156 nucleotides consisted of four segments, nt 1-37, 37-103, 103-140 and 140- 156 (Fig. 5D). Through analysis of the atomic distances between the atoms of Puf-A and 5.8S rRNA sequences, it was found that three segments of rRNA sequences from nt 37- 103, 103-140, and 140-156 intermingled directly with Puf-A protein (with atomic distances shorter than 5 Å). Since the orientation of the segment nt 1-37 is far flung from the Puf-A structure, it is suggested that this segment might not be involved in the binding with Puf-A (Fig. 5D). On the other hand, human Puf-A had an N-terminal PUF domain (blue), which consisted of tandem Puf repeats with an inserted non-Puf repeat region (pink) in the middle of the domain, and a C-terminal CPL domain (green). The molecular docking analysis revealed that the PUF domain recognized the segments nt 103-140 and 140-156 of 5.8S rRNA, whereas the CPL domain also interacted with the nt 37-103 (Fig. 5D). Such molecular interactions of Puf-A-5.8S rRNA complexes are further illustrated in Fig. S3c, where the concave surface of the PUF domain is the major area of Puf-A to recognize the double-stranded structures (nt 103-140) within the 5.8S sequence (Fig. S3c). This finding agrees with the previous report indicating that Puf-A could bind double-stranded nucleic acids (Qiu, McCann et al., 2014). Here, Puf-A is proposed to specifically interact with the 5.8S sequence within pre-rRNA in the nucleolus.

Since mature 5.8S rRNA resided within 60S ribosome in cytoplasm and the immature 5.8S (the 5.8S sequence within pre-rRNA) was present in pre-ribosomes in the nucleus (Thomson & Tollervey, 2010), Our finding of co-distribution and co-localization of Puf-A and 5.8S sequence in pre-rRNA (Fig. 5A and B) further confirm that Puf-A is associated with 90S pre-ribosomes within the nucleolus. Previously, mutation of yeast Puf6 (a homologue of mammalian Puf-A) led to reduction of mature 5.8S, suggesting its involvement in the processing of yeast 7S pre-rRNA (a precursor of 5.8S rRNA) to mature 5.8S rRNA (Qiu et al., 2014), thus facilitating the assembly of 60S ribosome, which presumably occurred in the cytoplasm (Li, Lee et al., 2009). However, in response to Puf-A silencing in various cells, steady state levels of mature 5.8S based on Northern blot analysis were not much altered (Fig. S3d), suggesting that Puf-A may not participate in the processing for the pre-5.8S rRNA.

Furthermore, silencing of Puf-A impaired the assembly of 90S in the nuclear fraction (Fig. 5E, left panel), resulting in the reduction of 80S ribosome and polysomes in the cytoplasm (right panel). Therefore, Puf-A interacts with the 5.8S region in the pre-rRNA, and is involved with the integrity of 90S pre-ribosomes.

### 7. Puf-A facilitates the assembly and nuclear export of pre-ribosomes

We further examined the subcellular localization of S6 and L5 protein in response to Puf-A expression. In MEFs, after transduction with c-Myc alone, the expression of S6 was mainly found in the cytoplasm (Fig. 6A, green, and 6B). By comparison, the expression of L5 was primarily in the nucleolus and also in the cytoplasm (red). When MEFs were co-transduced with Puf-A and c-Myc, the expression of S6 and L5 was obviously increased and both were co-localized in the cytoplasm (yellow). These results suggest that Puf-A promotes ribosome assembly, thereby facilitating maturation of the 80S ribosome.

**Figure 4.**
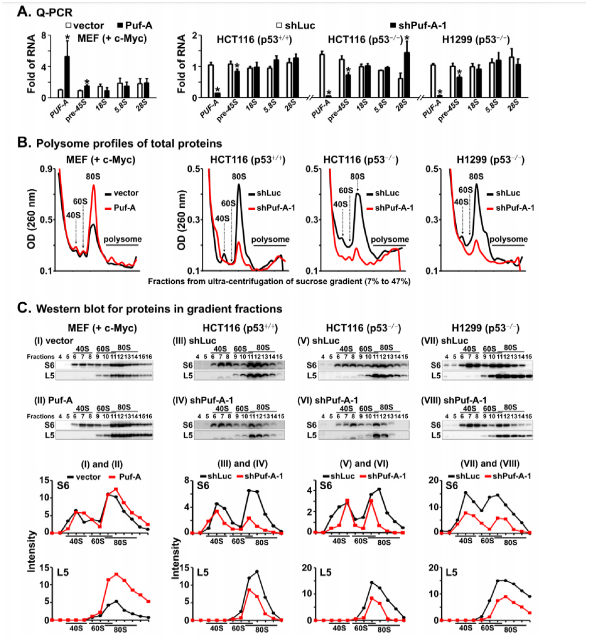
Role of Puf-A in ribosome biogenesis. **A**. Q-PCR analyses of the expression of *PUF-A*, pre-45S, 18S, 5.8S and 28S rRNAs in MEFs and cancer cell lines (p53^+/+^-HCT116, p53^-/-^-HCT116, and p53^-/-^-H1299), after various transductions were performed. GAPDH was used as an internal control. Mean ± SD values (n = 3) are shown. ^*^ refers to *p* < 0.05 (one-way ANOVA). **B**. Polysome profiles of ribosome extracts obtained from MEFs that either co-expressed Puf-A and c-Myc (red) or expressed c-Myc alone (black) were analyzed. Various fractions of ribosome extracts from 7 to 47% sucrose gradients after ultra-centrifugation were examined with ultraviolet light at 260 nm. Similarly, polysome profiles of ribosome extracts from cancer cell lines (p53^+/+^-HCT116, p53^-/-^-HCT116 and p53^-/-^-H1299) that were transduced with control shLuc (red) and shPuf-A-1 (black) lentivirus were analyzed. **C**. Equal volumes of protein extracts of various ribosomal fractions from MEFs, p53^+/+^- HCT116 cells and p53^-/-^-HCT116 and p53^-/-^-H1299 cells after transduction were precipitated using trichloroacetic acid and then analyzed via western blot. (I) and (II) represent protein compositions of various fractions after transduction of MEFs with vector and Puf-A; (III), (V), and (VII) with control shLuc; and (IV), (VI), and (VIII) with shPuf-A-1. The expression of S6 and L5 protein from MEFs, p53^+/+^-HCT116 cells, p53^-/-^- HCT116 and -H1299 cells after transduction was quantized by ImageJ software (Schneider, Rasband et al., 2012).

In various cancers, both L5 and S6 were co-localized in the cytoplasm (Fig. 6A, yellow) but rarely in the nucleolus in control cells, consistent with findings in MEFs with Puf-A and c-Myc co-expression. After silencing of Puf-A, the expression of S6 and L5 in these cells obviously decreased in the cytoplasm with increase in the nucleolus compared to control cells (Fig. 6A and 6B).

These results indicate that silencing of Puf-A leads to accumulation of ribosomal proteins in the nucleus, causing severe ribosome defects in cancer cells.

### 8. Puf-A recruits NPM1 in pre-ribosomes and prevents its translocation

To search for Puf-A interacting proteins that mediate the assembly and nuclear export of pre-ribosomes, we performed immunoprecipitation and mass spectrometric analyses which identified NPM1 was as a Puf-A-interacting protein in H1299 cells (Fig. S4a and b). Exogenous HA-tagged Puf-A co-precipitated with flag-tagged NPM1; endogenous Puf-A was also associated with NPM1 (Fig. 7A and B) in the nucleolus (Fig. 7C, shLuc). However, upon silencing of Puf-A, the NPM1 was translocated from pre-ribosomes in the nucleoli to the nucleoplasm of these cells (Fig. 7C, shPuf-A-1). In addition, silencing of Puf-A in cells did not alter the expression of NPM1 (Fig. 7D) compared to control cells. In cancer cells, NPM1 was found with 40S, 60S and 90S pre-ribosomes (Fig. 7E I, III and V). However, after transduction with shPuf-A-1, the expression of NPM1 in these pre-ribosomal fractions of nucleolus was decreased significantly, compared to control shLuc cells (Fig. 7E II, IV and VI). These results are consistent with the observation of NPM1 translocation in Fig. 7C.

**Figure 5.**
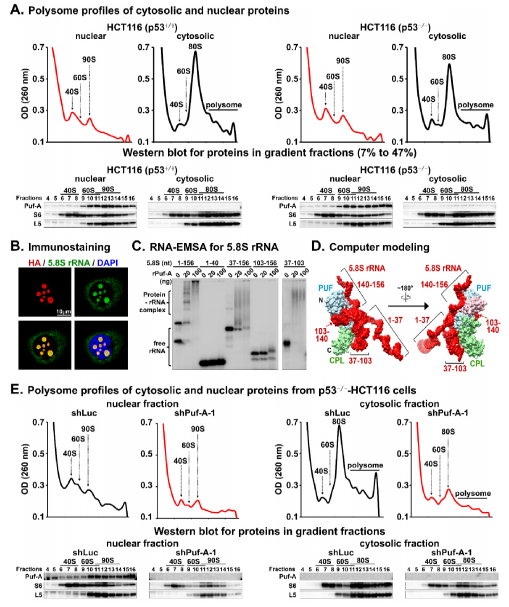
Interaction of Puf-A with pre-rRNA in 90S pre-ribosomes. **A**. Polysome profiles of cytosolic and nuclear extracts obtained from p53^+/+^- and p53^-/-^- HCT116 cells. Various fractions of ribosome extracts from 7 to 47% sucrose gradients after ultra-centrifugation were examined with ultraviolet light at 260 nm. Equal volumes of protein extracts of various ribosomal fractions were precipitated using trichloroacetic acid and then analyzed via western blot. **B**. Immunofluorescence staining of H1299 cells that overexpressed HA-tagged Puf-A with antibodies directed against HA and 5.8S rRNA sequence and counterstained with DAPI (blue). Specific interaction of HA-tagged Puf-A (red) with 5.8S sequence (green) within the nucleus (blue) is shown in yellow in the cells. **C**. RNA-EMSA analyses of rRNAs bound by rPuf-A *in vitro*. ^32^P internally labeled full-length 5.8S rRNA sequence (1-156 nt) and its truncated forms (nt 1-40, 37-156, 103-156, and 37-103) were prepared as described in Supplementary Information and incubated with increasing concentrations of rPuf-A. Autoradiographs of RNA-EMSA analyses are shown. The slowed motility corresponding to the protein-rRNA complex is indicated. **D**. Molecular interactions between Puf-A and 5.8S rRNA surface models. Structures of human Puf-A and 5.8S sequence within pre-rRNA were obtained from Protein Data Bank (PDB codes, 4WZR and 4V6X, respectively) (Khatter, Myasnikov et al., 2015, Qiu et al., 2014). Both Puf-A and 5.8S rRNA are shown here as solid surface models. The PUF domain (N131-E460) of Puf-A is marked in blue, whereas a non-PUF repeat sequence (E274-E349) is shown in pink; and the CPL domain (V461-L646) is shown in green. The molecular docking of the Puf-A protein-5.8S rRNA complex was performed with NPDock server(Tuszynska et al., 2015) as described in Materials and Methods. The molecular docking analysis indicates that the Puf-A interacts with the three segments (from nt 37 to156) of 5.8S rRNA, but not the nt 1-37. **E**. Polysome profiles of cytosolic and nuclear extracts obtained from p53^-/-^-HCT116 cells that were transduced with control shLuc and shPuf-A-1 lentivirus. Equal volumes of protein extracts of various ribosomal fractions were precipitated using trichloroacetic acid and then analyzed via western blot.

In MEFs after transduction with c-Myc alone, NPM1 was primarily expressed in non-ribosomal fractions, but not in pre-ribosomal fractions (Fig. 7E, VII). After co-transduction with Puf-A and c-Myc, there was an increase in the expression of NPM1 with the 40S, 60S and 90S pre-ribosomes in MEFs (Fig. 7E, VIII). These results suggest that Puf-A recruits NPM1 to pre-ribosomes and prevents its translocation from nucleolus to nucleoplasm, conceivably promoting the assembly of ribosomes.

**Figure 6.**
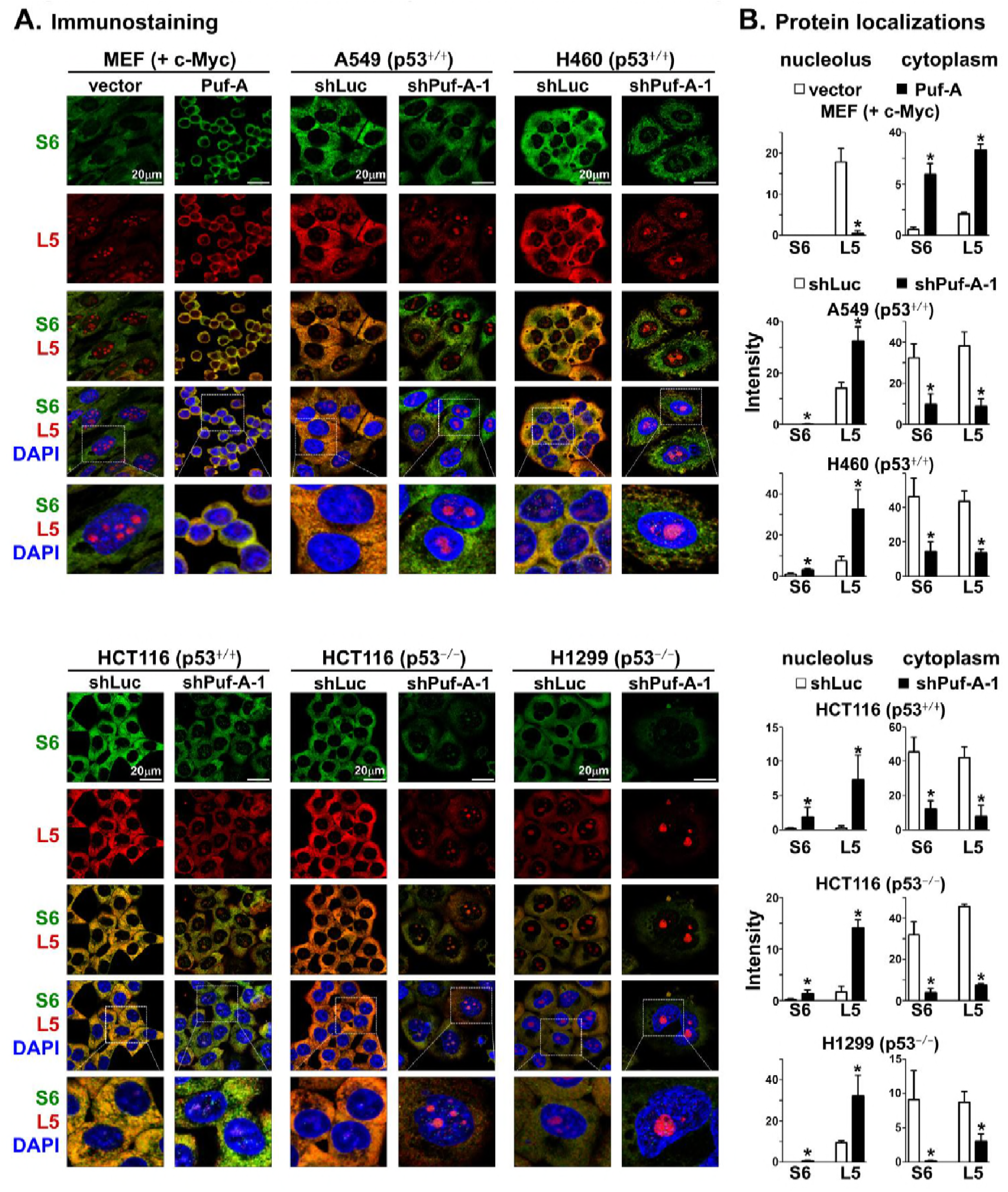
Ribosome assembly and nuclear export by Puf-A expression. A. Immunofluorescence staining of MEFs that overexpressed c-Myc alone or co-expressed Puf-A and c-Myc. Similarly, p53^+/+^-A549, -H460 and -HCT116 cells, as well as p53^-/-^-HCT116 and -H1299 cells, were transduced, separately, with shLuc (control) or shPuf-A-1 virus. Then, the cells were analyzed with antibodies against L5 (RPL5, red) and S6 (RPS6, green) and counterstained with DAPI (blue). Specific interaction between L5 and S6 in the cytoplasm is shown in yellow. Enlarged images of L5 and S6 staining in cells are shown in the lower panels. **B**. The expression of S6 and L5 proteins in the cytoplasm and nucleolus from MEFs, p53^+/+^-HCT116 cells and p53^-/-^-HCT116 and -H1299 cells after transduction was quantized by ImageJ software.

**Figure 7.**
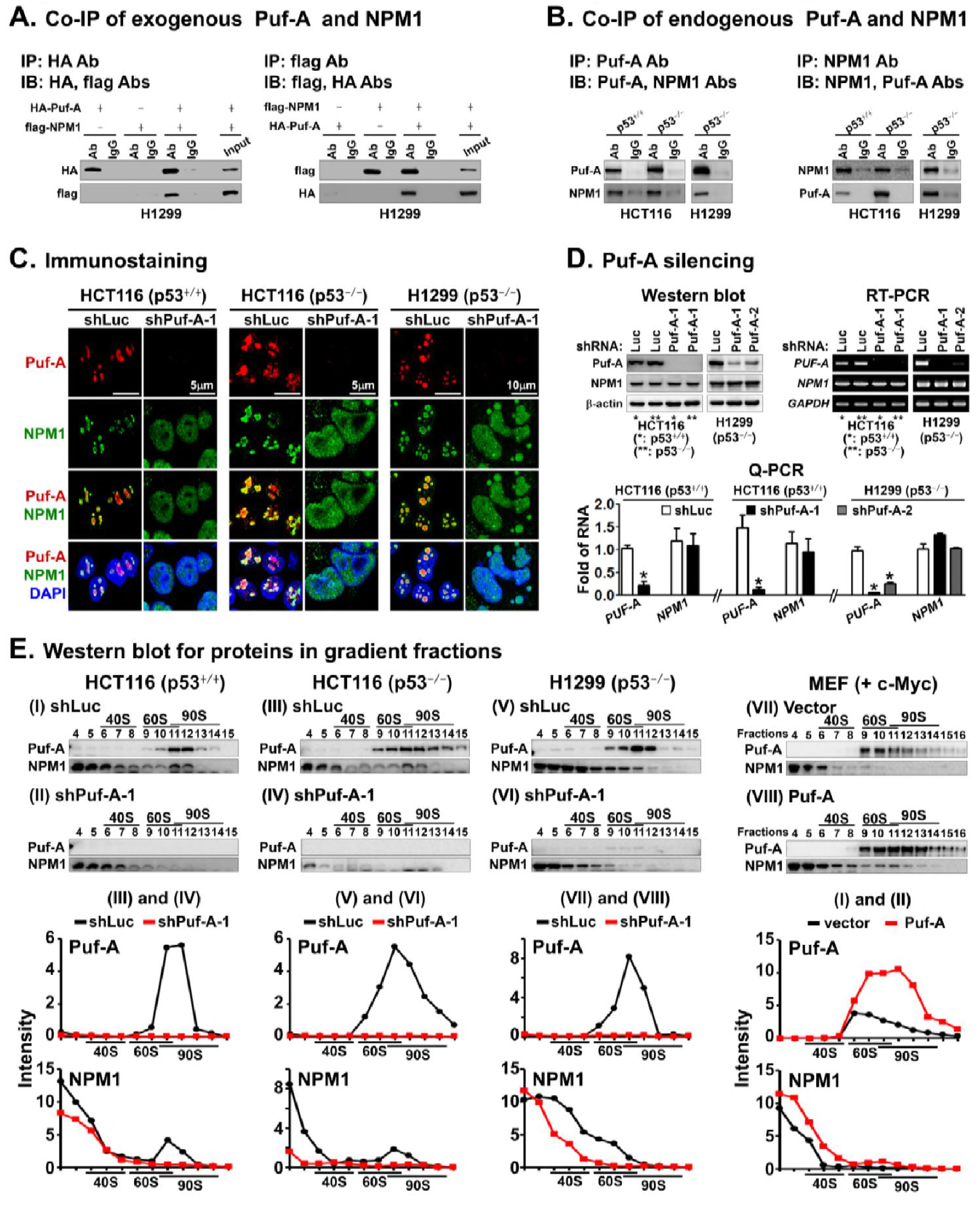
Induction of NPM1 translocation by Puf-A silencing. A. Co-immunoprecipitation (Co-IP) of exogenous Puf-A and NPM1 proteins. Cell lysates obtained from H1299 cells after co-transfection with HA tagged Puf-A (HA-Puf-A) and flag tagged NPM1 (flag-NPM1) overexpression vector, were co-IP with antibodies directed against HA or flag, and subsequently immunoblotted (IB) with tag antibodies as indicated. **B**. Co-IP of endogenous Puf-A and NPM1 proteins. Cell lysates from p53^+/+^HCT116, p53^-/-^HCT116 and H1299 cells were co-IP with antibodies directed against Puf-A or NPM1 and subsequently immunoblotted (IB) with Puf-A and NPM1 antibodies as indicated. **C**. Immunofluorescence staining for Puf-A and NPM1 expression in p53^+/+^- and p53^-/-^- HCT116 cells and p53^-/-^-H1299 cells. The cells were transduced with control shLuc and shPuf-A-1 and stained with antibodies directed against Puf-A (red) and NPM1 (green) and counterstained with DAPI (blue). **D**. Western blot, RT-PCR and Q-PCR analyses of the expression of Puf-A (*PUF-A*) and NPM1 (*NPM1*) in p53^+/+^-, p53^-/-^-HCT116 and p53^-/-^-H1299 cells after transduction with shPuf-A-1, shPuf-A-2 or control shLuc. β-actin and GAPDH were used as internal controls. In Q-PCR, mean ±SD values (*n* = 3) are shown. ^*^ refers to *p* < 0.05 (one-way ANOVA). **E**. Ribosome extracts obtained from p53^+/+^- and p53^-/-^-HCT116 and p53^-/-^-H1299 cells that were transduced with control shLuc and shPuf-A-1 lentivirus. Similarly, ribosome extracts obtained from MEFs that either co-expressed Puf-A and c-Myc or expressed c-Myc alone were also analyzed. Equal volumes of protein extracts in various ribosomal fractions from p53^+/+^- and p53^-/-^-HCT116, p53^-/-^-H1299 cells and MEFs after transduction were precipitated using trichloroacetic acid and then analyzed via western blot.

It was reported that NPM1 participates not only in the regulation of pre-45S rRNA transcription but also in the assembly and nuclear export of pre-ribosomes in various cancers (Maggi, Kuchenruether et al., 2008). Our results indicated that Puf-A may complex with NPM1 in nucleolus to mediate ribosome biogenesis.

## Discussion

As elaborated in graphic illustration (Fig. 8), we identified Puf-A as a critical factor for 90S pre-ribosome assembly, which regulates early ribosome biogenesis. In transformed and cancer cells, it was found that over-activation of ribosome biogenesis induced by Puf-A is required for Kras^G12D^/p53^-/-^–induced tumor progression and cancer growth, including tumors with *TP53* mutation. The over-activated ribosome biogenesis is consistent with the concept that cancer cells require extensive protein synthesis, which relies on a constant supply of new ribosomes (Dez & Tollervey, 2004).

**Figure 8.**
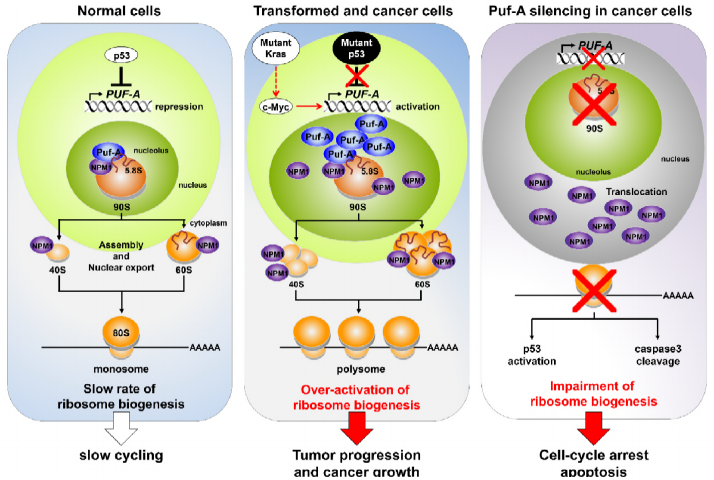
Graphical illustration.

Our clinical analysis of human lung cancer first demonstrated that the expression of Puf-A is upregulated in advanced human lung cancer and tumors, especially with *TP53* mutation, consistent with the finding that the expression of Puf-A in lung adenocarcinomas of Kras^G12D^/p53^-/-^ mice was higher than in Kras^G12D^/p53^+/+^ mice. More importantly, we are the first to demonstrate that the expression of Puf-A is highly prognostic for stage I lung cancer. To date, few markers have been proven to be predictive of the outcome of early stage lung cancer (Uramoto & Tanaka, 2014). Our new finding of Puf-A as prognostic marker for stage I lung cancer will address this unmet need in clinical practice.

Furthermore, Puf-A silencing alters the onset of adenomas and significantly prevents the progression to adenocarcinomas in the lungs of Kras^G12D^/p53^-/-^ mice. Silencing of Puf-A in various *TP53*–mutated cancers and c-Myc/p53^-/-^–transformed cells increased the expression of apoptotic proteins, such as cleaved caspase3 or cleaved PARP1, culminating in cell apoptosis.

Previously, higher expression levels of Puf-A was reported in human breast cancers with advanced clinical stages (Fan, Lee et al., 2013). However, the transcriptional regulation of Puf-A in cancer cells and its molecular mechanism underlying cancer promotion were unknown. Here, we have elucidated its key role in ribosome biogenesis, which contributed to tumor progression. In addition, we provided the first evidence that *PUF-A* locus harbors the c-Myc and p53 response elements, and the transcription of Puf- A is positively regulated by c-Myc and negatively regulated by p53. These finding were all summarized in graphical illustration (Fig.8).

In this study, we also elucidated several distinct features of Puf-A which set it apart from other PUF family proteins, including the differential roles of Puf6, a homolog of Puf-A in yeast, in ribosome biogenesis. In yeast, two assembly factors of 60S ribosome, Pwp1 and Nop12, were reported to participate in the folding of double-stranded structure of 5.8S sequence within pre-rRNA, thereby facilitating its processing (Talkish, Campbell et al., 2014).

We demonstrated that Puf-A specifically interacts with the nt 37-156 of 5.8S sequence within pre-rRNA of 90S pre-ribosome, but does not seem to be involved in its processing in the cytoplasm. First, the co-distribution of Puf-A with pre-rRNAs in polysome profiling of nuclear fraction and the co-localization of Puf-A and 5.8S sequence in pre-rRNA in immunostaining suggest that Puf-A might be associated with 90S pre-ribosomes in nucleolus. Secondly, steady state levels of mature 5.8S based on Northern blot analysis were not much altered in response to Puf-A silencing in cells. Thirdly, Puf-A expression seemed to involve in ribosome assembly and its nuclear exports. In other words, Puf-A might be important for proper folding of 5.8S sequence in pre-rRNA, thereby allowing various assembly factors to associate with 90S pre-ribosomes and facilitate ribosome biosynthesis. Thus, our finding regarding how Puf-A interacts with 5.8S sequence is in agreement with the proposal that Puf-A recognizes double-stranded nucleic acids (Qiu et al., 2014).

The classical PUF proteins (e.g., PUM1 or PUM2), which bind to specific recognition sequences in the 3’ untranslated regions (3’ UTRs) of mRNAs and control the stability and translation of transcripts, were found in the cytoplasm (Quenault et al., 2011). However, Puf-A was unique in its localization in the nucleolus and association with 90S pre-ribosomes in early ribosome biogenesis. In addition, previous report indicated that Puf-A could bind double-stranded nucleic acids without sequence specificity (Qiu et al., 2014), thus Puf-A may not participate in the processing of specific pre-rRNAs. Thus, Puf-A is distinct from its PUF homologs in its localization in the nucleolus and association with 90S pre-ribosomes.

As a result, Puf-A silencing induced the translocation of NPM1 from the nucleoli to the nucleoplasm, thus causing impairment of the assembly of 40S and 60S ribosomes and leading to defective synthesis of 80S ribosomes and polysomes (Fig.8). These findings are consistent with the notion that the translocation of NPM1 from nucleoli to the nucleoplasm occurs as a stress signal for ribosome biogenesis, resulting in growth inhibition of cancers (Lindstrom, 2011).

In contrast, overexpression of Puf-A in MEFs with c-Myc activation enhanced the activity of ribosome biogenesis and dramatically stimulated cell proliferation and transformation. It was found that Puf-A recruited NPM1 within 90S pre-ribosomes and facilitated the assembly of 40S and 60S ribosomes. Our results were consistent with previous studies indicating that NPM1, as a nucleolar–cytoplasmic shuttling protein, promotes the nuclear export of 40S and 60S ribosomes into the cytoplasm to increase the number of 80S ribosomes and polysomes, thus leading to increased cell proliferation (Maggi et al., 2008). It should be noted that there was no obvious alteration in proliferation rate or phenotypic conversion of normal MEFs in response to Puf-A silencing, suggesting that normal cells with low expression of Puf-A and slow rate of ribosome biogenesis might not be affected by Puf-A silencing.

Finally, our studies reveal that the expression of Puf-A links ribosome biogenesis to cancer progression and further provides a mechanistic explanation for Puf-A regulation of 90S pre-ribosome assembly. The anti-cancer effects of targeting Puf-A were observed in many cancers, including lung, colon, and breast cancers. Thus, the study provides a strategy to treat a variety of cancers.

## Materials and Methods

Experimental procedures are detailed in the Supplementary Information.

## Acknowledgments

This study was supported by grants (NSC 102-2321-B-182A-004, MOST 103-2321-B- 182A-003 to MOST 104-2321-B-182A-001 and MOST 106-3114-B-182A-001) from the Ministry of Science and Technology of Taiwan and (OMRPG3C0041 to OMRPG3C0044) from CGMH at Linko of Taiwan to J.Y.

